# DeepLoco: Fast 3D Localization Microscopy Using Neural Networks

**DOI:** 10.1101/267096

**Authors:** Nicholas Boyd, Eric Jonas, Hazen Babcock, Benjamin Recht

## Abstract

Single-molecule localization super-resolution microscopy (SMLM) techniques like STORM and PALM have transformed cellular microscopy by substantially increasing spatial resolution. In this paper we introduce a new algorithm for a critical part of the SMLM process: estimating the number and locations of the fluorophores in a single frame. Our algorithm can analyze a 20000-frame experimental 3D SMLM dataset in about one second — substantially faster than real-time and existing algorithms. Our approach is straightforward but very different from existing algorithms: we train a neural network to minimize the Bayes’ risk under a generative model for single SMLM frames. The neural network maps a frame directly to a collection of fluorophore locations, which we compare to the ground truth using a novel loss function. While training the neural network takes several hours, it only has to be done once for a given experimental setup. After training, localizing fluorophores in new images is extremely fast — orders of magnitude faster than existing algorithms. Faster recovery opens the door to real-time calibration and accelerated acquisition, and future work could tackle more complicated optical systems and more realistic simulators.

## 1 Introduction

Visualizing microscopic biological processes is crucial to understanding their function; optical microscopy has been a major tool of biological investigation for over a hundred years. Over the past decade, fundamental physical limits in the resolution of classical microscopy systems have been surmounted by superresolution microscopy techniques [36, 39], enabling the visualization of cellular structures far smaller than before. 3D single-molecule localization microscopy (SMLM) localizes individual fluorophores in 3D space in order to generate an image [11, 20] or facilitate analysis of fluorophore locations. However, these techniques come at a computational cost, requiring advanced algorithms to perform the reconstruction.

While the ultimate goal of most SMLM experiments is to generate high-resolution images, the process involves several intermediate steps with different products. A SMLM experiment proceeds in four stages. First, fluorophores, each a few nanometers in size, are attached to the sample and then stimulated to stochastically fluoresce. Second, a sequence of frames are taken using an optical microscope. Due to the stochastic stimulation, only a small subset of fluorophores are active in each frame. In the third stage, each frame is analyzed to determine the location of each fluorophore active in the frame. (A more sophisticated method processes multiple frames together.) Finally the collection of all locations from all frames is analyzed directly, or used to render high-resolution image in two or three dimensions.

The algorithms we develop in this paper address the third step of a SMLM experiment, analyzing a single frame to produce a short list of locations of the fluorophores active in the frame. While the aim of most SMLM experiments is to produce an image, we will refer to the task of localizing the fluorophores in a single frame as the SMLM inverse problem.

The localization microscopy community has yet to agree on a universal metric by which to measure performance [37]. Several proposed quality metrics operate on the estimated locations and number of fluorophores directly (such as the Jaccard index at a particular radius), while others apply to the final product of a rendered, high-resolution image (e.g., PSNR). In this paper we propose, and directly optimize, a new kind of metric: the mean squared error between an infinite-resolution image generated from the estimated fluorophores and the image generated by the ground truth fluorophores. While this metric is an image-based distance, it operates directly on sets of fluorophores.

Our approach to the SMLM inverse problem is to harness the availability of fast, accurate forward-model and noise simulators to train a neural network that maps a single frame to a list of localizations. We do this by attempting to minimize expected loss on simulated data. In the language of statistics, we are attempting to approximate the Bayes’ estimator with a neural network. Compared to traditional maximum-likelihood algorithms, our method is easier to calibrate, orders of magnitude faster (once trained), works with a wider variety of noise and forward models, and achieves equal performance on several 2D and 3D datasets.

Our method requires an accurate simulator. Unlike traditional maximum-likelihood/convex optimization based approaches, which make strong assumptions about the forward and noise models (specifically that the log-likelihood is concave and that the forward model is linear), our method can be applied to problems with arbitrary noise statistics, aberrations, and non-linear forward models. Furthermore, we do not require a functional form for the forward simulator, which allows us to handle non-deterministic forward models that take into account aberrations such as dipole effects [11] and, perhaps more importantly, allows us to generate training data directly from a few Z-stacks.

We list three possible disadvantages to our approach. The first is that training a neural network is, at the present time, difficult: unlike convex optimization, there are a plethora of hyperparamters and training usually requires a human in the loop. The second disadvantage is that the method requires an accurate end-to-end generative model that includes the variations and aberrations that will be encountered in the real experimental setup — though arguably this is advantageous: maximum-likelihood approaches are unable to take advantage of this kind of prior knowledge. As we describe in §5, the generative model we use is extremely simple and requires only a single Z-stack of experimental data.

Finally, the third disadvantage is common to all applications of neural networks: there is, at present, essentially no theoretical understanding or performance guarantees. For example, training could, in principle, fail for a new experimental setup. Additionally, any given reconstruction could fail. While this is, indeed, a potential shortcoming, experimental evidence suggests that our method is (in practice) at least as reliable as convex methods. Furthermore, all theoretical performance guarantees for convex optimization/maximum likelihood based methods rely on very strong assumptions about the data generation process. We remind the reader that if these assumptions are violated (and they often are, in practice) the conclusions of the theory do not apply. That said, maximum-likelihood based approaches are used extensively in applications where the theoretical assumptions are violated and have a very long history of reliability.

The paper is organized as follows. First, §2 introduces common notation. In §3 and §4 we describe two bodies of related work, provide background material for our approach, and put our method in context. In §5 we give details on our approach, and in §6 we describe our experimental setup and results. Finally in §7 we describe several possible extensions of our method; some simple, others speculative.

While preparing this manuscript, we discovered another paper that applies deep learning techniques to STORM microscopy [29]. The major difference between our approaches is that while our algorithm returns a set of localizations, DeepStorm returns a single gridded image. This choice limits the approach to 2D (it’s unclear that dense reconstruction could be extended to 3D without, at the very least, a huge increase in computation time), limits rendering to a single scale, and precludes any downstream analysis of the fluorophore locations [7, 10, 25, 30]. Furthermore, the use of an ℓ_1_ penalty to encourage sparsity in the reconstructed image introduces an additional parameter that must be tuned and prevents interpretation of the algorithm as an approximation of the Bayes’ estimator. With that said, there are some interesting similarities between the approaches: the loss function is essentially a gridded version of our loss function, and both algorithms are substantially faster than existing algorithms. Unfortunately we are unable to directly compare the two approaches as the code for [29] is not yet available.

## 2 Notation and Loss Functional

One issue with SMLM as an inverse problem is the choice of loss function. In many inverse problems the object to be estimated is an element of a finite-dimensional Hilbert space, in which case the *L*_2_ distance is a natural loss function. SMLM is more complicated: it is unclear how to compare two sets of points. In this section we argue that, in fact, SMLM is not so different from simpler inverse problems: while the intermediate output of a SMLM experiment may be a collection (of varying cardinality) of (possibly weighted) point sources, the final objective of most SMLM experiments is to render an image. This interpretation suggests a natural metric: the squared error of the resulting rendered image. We propose rendering the image at *infinite* resolution for computational efficiency and using the resulting *L*_2_ distance directly as the training loss function.

In this section we introduce some common notation for the remainder of the paper and formalize the loss function described above. The underlying object to be estimated is a set of points in Θ ⊂ **R**^2^ (or Θ ⊂ **R**^3^ for 3D SMLM), *γ* = {*θ*_1_,…, *θ*_n_}. Here *θ*_*i*_ ∈ Θ is the location of the *i*-th fluorophore in space. Note that *n*, the number of fluorophores active in the frame, is unknown and varies from frame to frame. While the object to be estimated is simply a collection of points, we’ll often deal with *weighted* sets of points, of the form {(*w*_1_, *θ*_1_),…,(*w*_*n*_, *θ*_*n*_)} where *w*_*i*_ ∈ **R** and *θ*_*i*_ ∈ Θ. The *w*_*i*_ will have different interpretations in different contexts. When we talk about simulating data or using maximum-likelihood estimation, *w*_*i*_ > 0 will be the intensity of the *i*-th active fluorophore in the frame — that is, roughly proportional to how many photons that fluorophore emitted during the exposure. In the output of the neural network, however, *w*_*i*_ will be interpreted as a confidence. We’ll often talk about unweighted sets of points (like γ) as weighted collections, in which case we’ll take each *w*_*i*_ to be one. Finally, it’ll often be convenient to make use of a bijection between weighted collections of (unique) objects in Θ and *finitely-supported* atomic measures on Θ. The bijection associates a weighted collection *γ* = {(*w*_1_, *θ*_1_),…, (*w*_*n*_, *θ*_n_)} with the measure

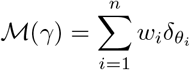

Here *δ*_*θ*_ is a point-mass supported on *θ*. Similarly, if *μ* is a finitely-supported atomic measure on Θ, 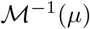 is a well-defined weighted set of points.

## 2.1 Rendering and loss functional

The final stage in many SMLM experiments is to render an image from the localized fluorophores. In practice, localizations must be convolved with a small convolution kernel before they are rendered. This blur serves two purposes: first, it allows an image to be formed, and second it attempts to make explicit uncertainty in the localization, both from the estimation process and from the fact that the fluorophore molecules (which are attached to the molecules of interest) have nonzero spatial extent. While in many SMLM applications the resulting images are rendered on a fine grid, we will simply consider the infinite-resolution image as a functional on **R**^2^ or **R**^3^. Given a convolution kernel *ϕ* ∈ *L*^2^(**R**^2^) (or **R**^3^), the image generated by the (confidence-weighted) set 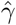 is

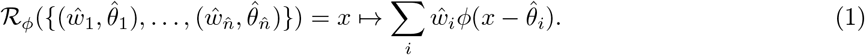

ℛ_*ϕ*_ can also be written compactly as a convolution of the measure 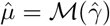 with the kernel *ϕ*:

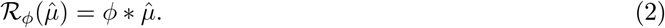

With this notation in place, we can introduce the family of loss functions we will use. With convolution kernel *ϕ*, let

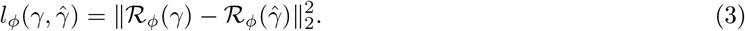

This expands into the following quadratic form in (*w*, 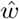):

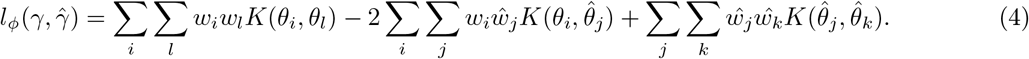

In the above *K*(*θ*, *ζ*) is the positive semi-definite function defined by

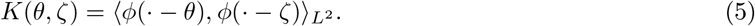

If *γ* is the true collection of fluorophore locations, we take each *w*_*i*_ to be identically one, while a localization algorithm may use 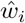 to encode its confidence that there is a fluorophore at location 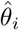.

As long as *K* is known, we can compute (3) efficiently, at least when *n* and 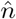 are relatively small. If *n* or 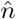 are large, any number of truncated or random embeddings will work to approximate (3) [8, 34, 43].

For instance, a typical choice of *ϕ* in applications is the standard Gaussian probability density function at a particular scale *σ*:

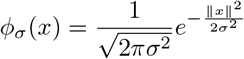

which corresponds to

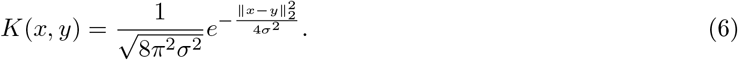

As we’ll see later in §5, more exotic choices are possible. In practice, *σ* is often chosen to be near the expected localization precision of the system, i.e., 20 to 50 nanometers.

## 3 Maximum-likelihood Methods

In this section we briefly describe existing techniques, almost all of which are based on maximum-likelihood estimation. Maximum-likelihood and regularized maximum-likelihood methods for inverse problems have proven to be effective over a wide variety of applications, and SMLM is no exception: the highest-performing SMLM algorithms are all variations on maximum-likelihood estimation [18, 38]. In this section we describe one family of convex approximations to the SMLM maximum-likelihood estimation problem.

These approaches assume additional structure in the measurement process and the noise model, though they seem to work well even when the assumptions aren’t met. First, they assume that the measurement process is a function of (only) the positions and intensities of the sources and is given by an operator 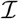. Furthermore, they require that 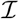 is additive in the sources and linear in the intensities:

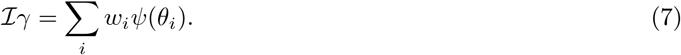

In the above, *ψ*: Θ → **R**^*d*^ is a known function. In microscopy, *ψ* is a spatially-translated and pixelated copy of the microscope’s on-axis point-spread function. These algorithms further assume that the negative log-likelihood of the noise distribution is a known convex function ℓ: **R**^*d*^ × **R**^*d*^ → **R**. For instance, if the per-pixel noise is approximately Gaussian, then 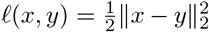 A maximum-likelihood estimate of γ is a solution to the following optimization problem:

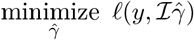

Even with these additional assumptions, this optimization problem is quite difficult: 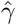 is of unknown cardinality, and the objective function is non-convex in the spatial locations of the fluorophores.

One way to avoid these issues is to lift the optimization variable 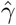 to the measure 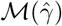. The additional structure described in (7) means that the nonlinear measurement operator 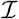 can be extended to a *linear* operator on measures. For instance, with 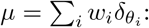

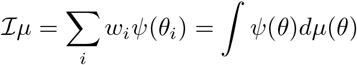

This last expression is well-defined for *all* signed measures of finite total variation. As the composition of a linear operator and a convex function is convex, the following optimization problem is convex in the variable **μ**:

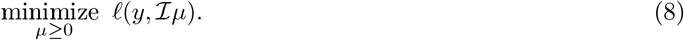

Unfortunately, the solution to (8) is, in general, not finitely-supported and thus cannot be interpreted as a weighted collection of points. One heuristic to encourage the solution of (8) to be supported on a small number of points is to add a penalty term on the total mass of **μ**: this is the infinite-dimensional analog of the ℓ_1_ norm. This modification results in the following (infinite-dimensional) convex optimization problem:

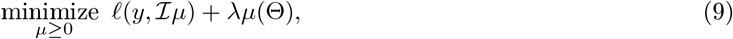

where *λ* is a positive parameter. It can be shown that the solution to (9) is guaranteed to be finitely supported [3, 5], and thus can be interpreted as a weighted collection; in practice, the support of the solution is often extremely sparse. Here *λ* > 0 allows us to trade off data fidelity for the cardinality of the support of the estimated measure. State-of-the-art algorithms for SMLM solve (9)[3] or a finite-dimensional, gridded analogue of (9)[14, 28, 32, 51].

In practice, these algorithms may also require postprocessing: for instance thresholding by removing points for which the estimated intensity 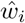 is low, or by clustering nearby localizations [45]. Of some interest are the myriad of theoretical results concerning (9): these results stipulate that if the measurement model 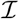 is accurate and some additional technical assumptions are satisfied, the solution to (9) is guaranteed to be close (in some sense) to the ground truth [13, 40].

## 4 Function Approximation Methods for Inverse Problems

In this section we briefly discuss related work applying deep learning techniques to inverse problems.

Recently there has been great interest in using deep neural networks for inverse problems, especially problems in imaging [26, 27]. We group the field into into two broad groups: amortized or compiled inference, and iterative, or unrolling approaches.

Amortized or compiled inference attempt to directly learn an approximation 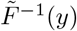 1 for the problem *y* ≃ *F(x)*, often by training a network with a very large number of known (*x*, *y*) pairs. In the applications highlighted in [26], including superresolution imaging [22], motion deblurring [49], and denoising [50], the output of 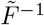 is a dense image. Other recent work [21] has attempted to learn an inverse model to classify the hand-written MNIST dataset from a camera system with minimal optics. In this case the predicted output of the network is an integer from zero to nine. Learning function approximators for inverse problems has a rich history, and multilayer neural networks were used beginning 30 years ago for inverse problems [23], including problems in optics [46]. More contemporary work on *compiled inference* for probabilistic programming [24] also extends this approach.

Unrolling approaches take an existing iterative algorithm and replace some components with a learned operator — either the actual iterative steps themselves (hence the “unrolling”) or a proximal operator for algorithms with a proximal step. These approaches can exploit known linearities in the problem. One of the earliest examples is learned iterative shrinkage-thresholding algorithm (LISTA [15]), which unrolled an iterative-shrinkage and thresholding algorithm and learned approximations for the adjoint and gram matrix. ADMM-based approaches [47] learn networks that approximate subproblems in ADMM.

In microscopy, recent work has attempted to learn data-driven methods for upsampling conventionally acquired images [35, 48], although in all cases, these data-driven methods make assumptions about the nature of the system under study. Indeed, the authors of [48] specifically caution against using their method for imaging novel biological structures.

## 5 DeepLoco

In this section we describe how we solve the SMLM inverse problem using a function approximator: we train a neural network to directly minimize expected per-frame loss on simulated data. We first describe the generative model we use to create training data. We then describe the loss function and how it can be extended to arbitrary problems involving weighted collections of objects. Finally we briefly describe the architecture of our network and how we train it.

### 5.1 Data generation

The success of function approximation techniques on novel datasets relies on the availability of vast quantities of labeled training data. We argue that SMLM falls into this class of problems. The combination of a reasonable generative model for fluorophores and a well-understood forward model means that, considered as a machine learning problem, SMLM has essentially *infinite* training data: we can simulate as many training examples as we need. Obviously there is still mismatch between the simulated data and any test data we encounter in the real world; the real question is how this mismatch affects the performance of our algorithms. In this paper we show that the mismatch is small enough that we can obtain good localizations.

The first step in simulating SMLM data is to generate random collections of fluorophores. For each image we sample the number of fluorophores, *n*, from a uniform distribution. We then sample *n* spatial locations independently and uniformly from a 3D box. We sample the fluorophore intensities from a uniform distribution.

Next, we run the collection of fluorophores through a forward model to generate noiseless observations. The forward model we use can be thought of as aggressive data augmentation and is very easy to apply in practice. We use laterally translated — in most experimental setups the PSFs are invariant to translation in X and Y — versions of an empirically measured PSF to generate new data. This approach has several advantages over using a fitted functional form. By taking multiple Z-stacks of different fluorophores (or beads), we can train the network to be robust to aberrations that vary from fluorophore to fluorophore (such as dipole effects [11]). It also removes the critical preprocessing step of fitting a parametric model to the Z-stacks and is hyperparameter-free.

The final step in simulating SMLM data is to add noise to the image. In our experiments we simply use Poisson noise for each pixel. While this is not a great fit for experimental data, we find that the method is robust enough that this mismatch is not an issue. As we discuss in §7, including a more accurate noise or background model could improve the performance of our approach.

### 5.2 Loss functions

In §2 we introduced a metric for SMLM. While that metric can be used to evaluate the results of a entire SMLM experiment, in this subsection we show how we use it on *single* frames in order to train a neural network. We also describe practical extensions, such as rendering at multiple scales to help training.

To compute the loss on a single frame we set the weights for the true active fluorophores to one, resulting in the target collection γ = {(1, *θ*_1_),…, (1, *θ*_*n*_)}. As the number of fluorophores active in a single image is relatively small (in our experiments at most a few hundred), we use (5) to compute *l*_*ϕ*_. For training we use the Laplacian kernel:

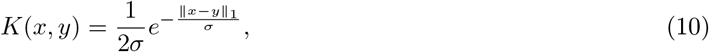

which corresponds to using the first modified Bessel function of the second kind as the convolution kernel *ϕ*.

In our experiments we found that evaluating the loss function at multiple scales during training improves the final performance of the network. The loss function is then

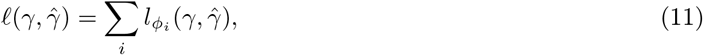

where each *ϕ*_*i*_ is a convolution kernel at a different scale. This loss function can be evaluated using the quadratic form in (5) with *K*(*x*,*y*) = Σ_i_*K*_i_(*x*, *y*), where 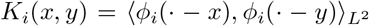 and each *ϕ*_i_ convolution kernel at scale σ*i*.

### 5.3 Generalization to other machine learning problems

Readers familiar with reproducing kernel Hilbert spaces [1] will recognize (10) and (6) as reproducing kernels. Indeed, another way to think of the metric (5) is as the *maximum mean discrepancy* [16] between the measures 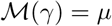 and 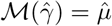 The maximum mean discrepancy between *μ* and 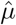 is given by

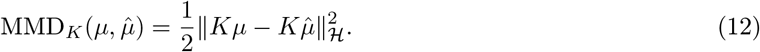

**Figure. 1.**
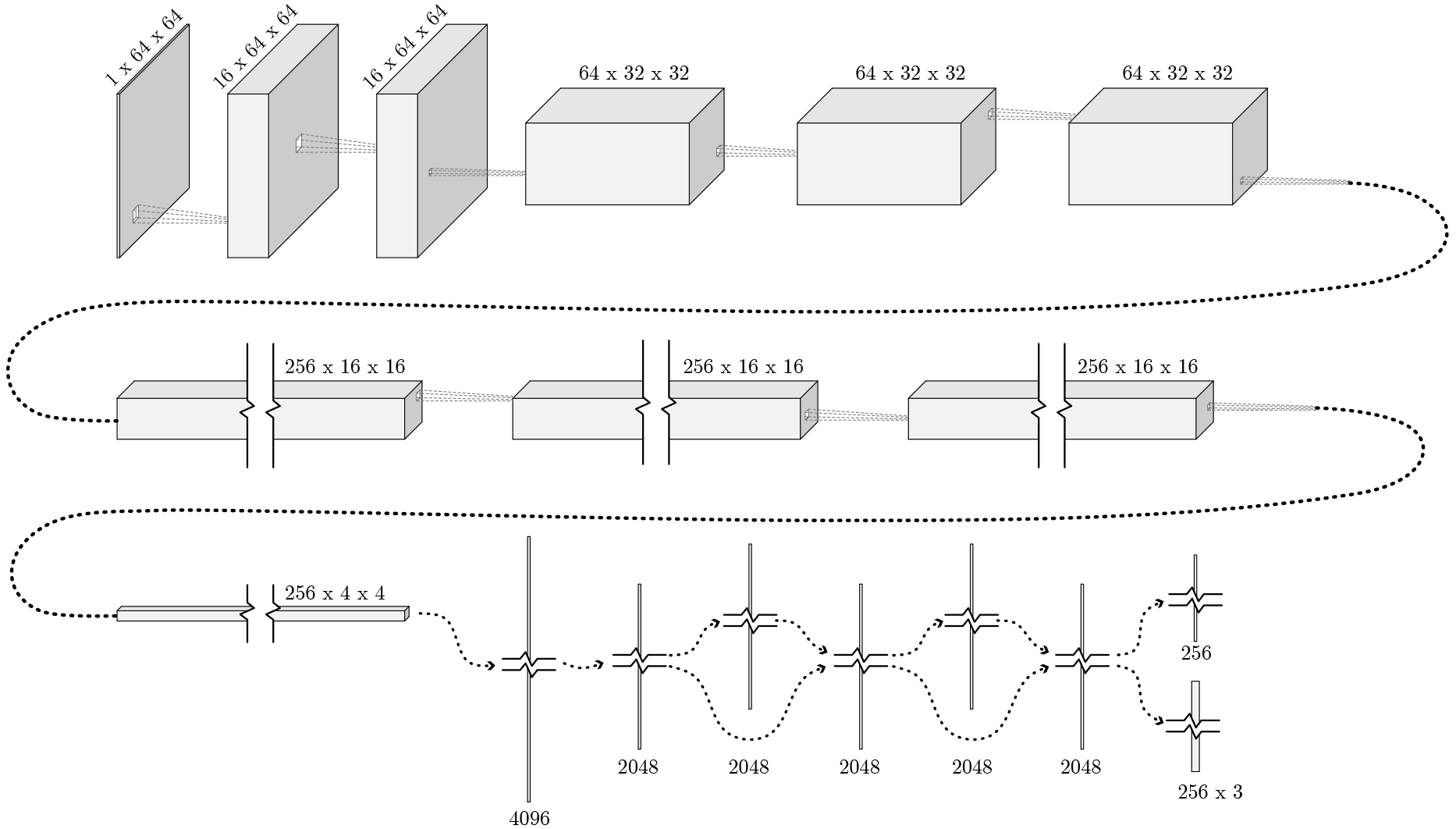
A visualization of the DeepLoco architecture. The first two convolutions use 5×5 filters, while the remaining convolutions use 3×3 filters. Spatial downsampling is by strided convolution; twice using 2 × 2 filters with stride 2 and once with 4×4 filters with stride 4.

In the above, 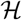 is the RKHS generated by the kernel *K*. By a slight abuse of notation we use *K* to denote the linear operator (with codomain 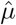) defined by

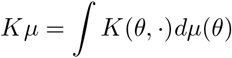

This suggests an extension of the loss function (3) to arbitrary spaces Θ equipped with a kernel *K*: simply take

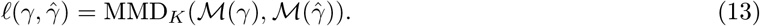

For more on the topic of embeddings of weighted collections of points (and general measures) into RKHS, see [42].

### 5.4 Neural network architecture

We use a fairly standard convolutional neural network architecture. We emphasize that our contributions are the application of neural networks to the localization problem and the loss function described above: the architecture of the neural network we use is essentially arbitrary and almost surely suboptimal. We use the same architecture for both 2D and 3D experiments (except for the final layer, which outputs either two or three spatial coordinates). The first part of the network is fully convolutional (with so-called ReLU nonlinearities) and alternates three times between performing convolutions at given scale and spatial downsampling by strided convolution. This is followed by a two-layer fully-connected ResNet [17]. Finally, the output of the ResNet is fed into two linear layers that output a fixed (but large) number, *K*, of sources. One linear layer outputs *K* weights; non-negativity of the weights is enforced by a ReLU nonlinearity. We take K to be much larger than *n*, the number of true sources; the network is free to set many of the weights to zero. The second linear layer outputs a tensor of size *K* by 2 (or 3, for 3D localization) that encodes the
predicted spatial locations 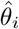. This layer uses a sigmoid nonlinearity to ensure that the estimated locations remain within a given spatial extent.

### 5.5 Training

We first simulate a batch of data to use as a validation set during training. During each training iteration, we simulate a new batch of training data (both spatial locations and noisy images) and run one step of a stochastic gradient descent variant (in our experiments either SGD with momentum or ADAM). Every few iterations we evaluate the error on the validation set; when the error plateaus we reduce the stepsize and reset the optimization algorithm.

## 6 Experimental Results

In this section we compare our algorithm to the existing state-of-the-art algorithms on both simulated and contest data. The SMLM community has established several [38] contests to objectively benchmark the performance of these algorithms for both 2D and 3D localization. In both cases, we compare to the best performing algorithms for each task: the Alternating Descent Conditional Gradient method (ADCG) [3] which won the 2016 high-density 2D challenge, and Spliner [2] (also referred to as CSpline), the winner (for the astigmatic PSF in low density and the double-helix PSF in both high and low densities) of 2016 3D challenge. Note that for data generated from our simulator (figure **??**) we give both competing algorithms a handicap: we run them across a range of parameter settings and post-hoc pick the one that gives the highest Jaccard index. While not feasible in the real world, this helps lessen the risk of “parameter-hacking”2

To compare the different algorithms in different regimes we vary both the source density (number of simultaneously active sources per frame) and the signal-to-noise ratio of the point sources. Reconstruction accuracy can be measured by comparing the estimated locations directly to the true locations or by comparing high (or infinite) resolution rendered images; we compute metrics of both types. In all experiments that compare localizations directly we use simple postprocessing on the output of our method: we cluster the output points into connected components in a thresholded distance graph and then remove points with low confidences. To compare localizations directly we follow [38] by first solving an assignment problem in euclidean space: matching each detected point to a nearby true source point in a manner that minimizes the total euclidean distance between pairs of points. We then consider all pairs of matched points closer than a threshold (in our experiments, either 50nm or 100nm) as true positives (TP), and all other source points as false positives (FP). Similarly, missed ground truth points are considered false negatives (FN). We then compute the Jaccard index, J,

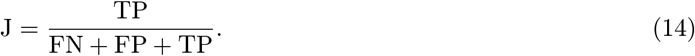

A Jaccard index of 1.0 indicates a perfect matching — all source points are recovered within the tolerance radius, with no spurious detections. With a fixed point matching we also compute the mean distance between matched pairs, in either *x* (2D) or *x* and *z* (3D). Finally, we report the image loss defined in (3) for a Gaussian convolution kernel with σ = 50nm.

### 6.1 2D SMLM

We first compare ADCG and DeepLoco on synthetic data to investigate how each algorithm performs under varying source conditions. The two dimensional synthetic data is generated from the 2016 SMLM 2D contest Z-stack. Fluorophores and noisy images are generated within 350nm of the focal plane over a 6.4*μ*m × 6.4*μ*m area with a per-pixel resolution of 100nm. We present the results in Figure 2. DeepLoco and ADCG have similar recognition accuracy (Jaccard index) and spatial localization accuracy across a variety of point densities and source intensities.

We also compare ADCG and DeepLoco on the MT0.N1.LD and MT0.N1.HD datasets from the 2016 SMLM contest in Table 2. We find that DeepLoco comparably if not slightly better than ADCG in the low and high-density cases. This is likely due to the relatively simplistic Gaussian PSF model used by ADCG, which is a poor fit for some of the datasets’ point sources which do not lie close to the focal plane.

**Figure. 2.**
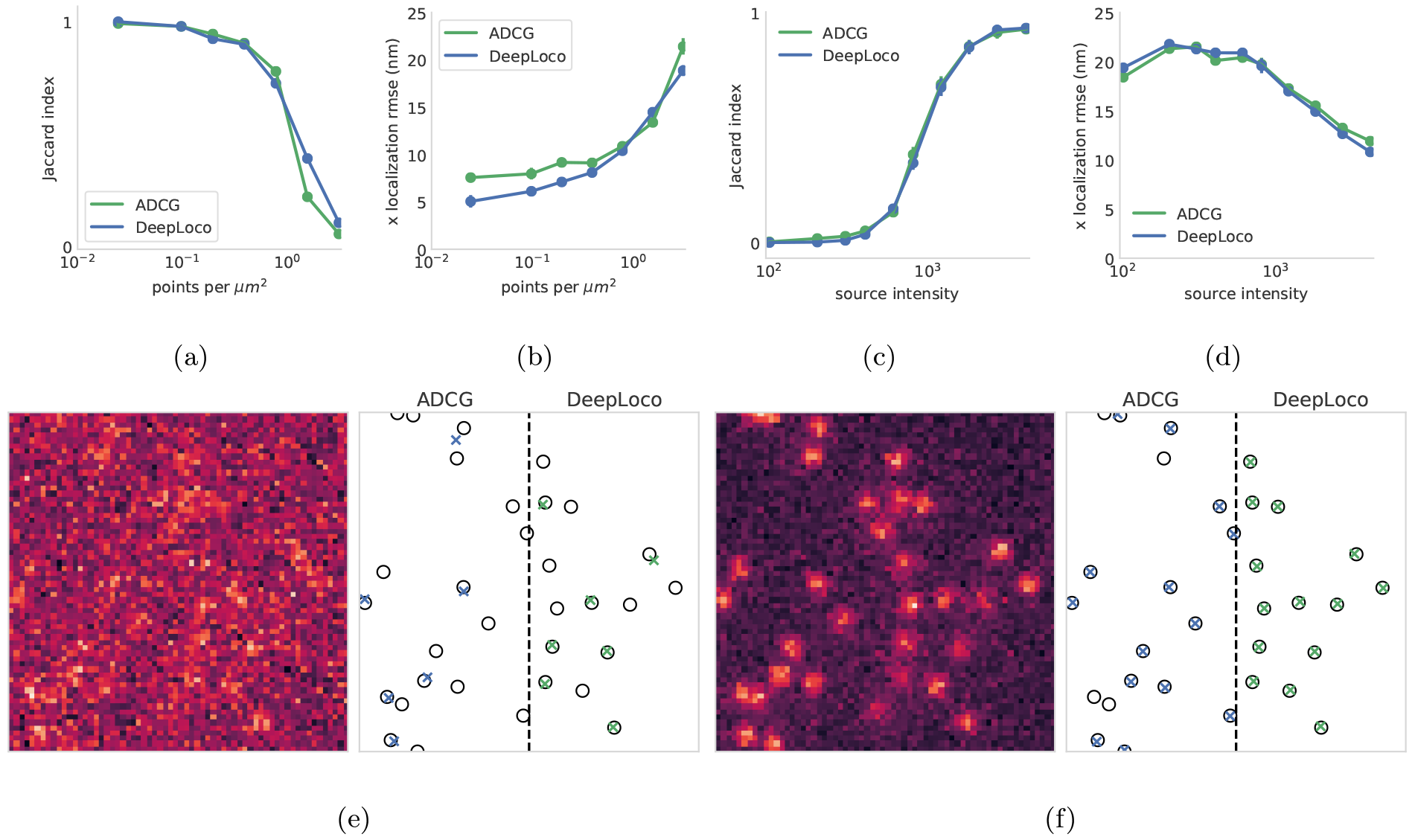
2D localization using DeepLoco and ADCG. All point matchings are computed with a 50nm tolerance radius. We vary the source density (**a.** and **b.**) and source intensity (**c.** and **d.**) and compare localization performance via Jaccard index and RMSE in the *x*-coordinate. For experiments with varying source intensity we use a fixed density of 0.2 molecules per *μ*m^2^. We find DeepLoco performs virtually identically to ADCG across the full range of tested parameters. **e.** and **f.** show example source images and the localizations returned by each algorithm.

### 6.2 3D SMLM

Three-dimensional SMLM uses point spread functions that vary with source depth, allowing a fluorophore’s position to be estimated in all three coordinates from a single frame. We compare DeepLoco to Spliner with two different PSFs: an astigmatic PSF [19] and a double-helix PSF [33].

We first compare the algorithms on synthetic data generated from the SMLM challenge calibration Z-stacks. These experiments (in Figure 3) show that DeepLoco significantly outperforms Spliner in terms of Jaccard index, while Spliner is slightly better in localization accuracy with the astigmatic PSF.

We next compare the algorithms on the MT0.N1.LD and MT0.N1.HD datasets from the 2016 SMLM challenge. We evaluate the results visually in Figure 4 and using quantitative metrics in Table 1. DeepLoco significantly outperforms Spliner in terms of Jaccard index and the kernel loss. The two algorithms are comparable in terms of spatial localization accuracy, except for the low-density double-helix data, where Spliner significantly outperforms DeepLoco.

### 6.3 Runtime

DeepLoco is significantly faster than ADCG and Spliner. While DeepLoco runs in (essentially) constant time regardless of fluorophore density, both ADCG and Spliner have iterative components that scale with the number of input sources. We run the algorithms on subsets of data from the SMLM 2016 contest to capture the runtime dependence on source density (Table 3). DeepLoco was run on an Amazon Web Services p3.2xlarge instance with a single Nvidia V100 GPU. ADCG and Spliner were run on the equivalent of an Amazon Web Services c4.8xlarge machine with 18 physical cores.

**Figure. 3.**
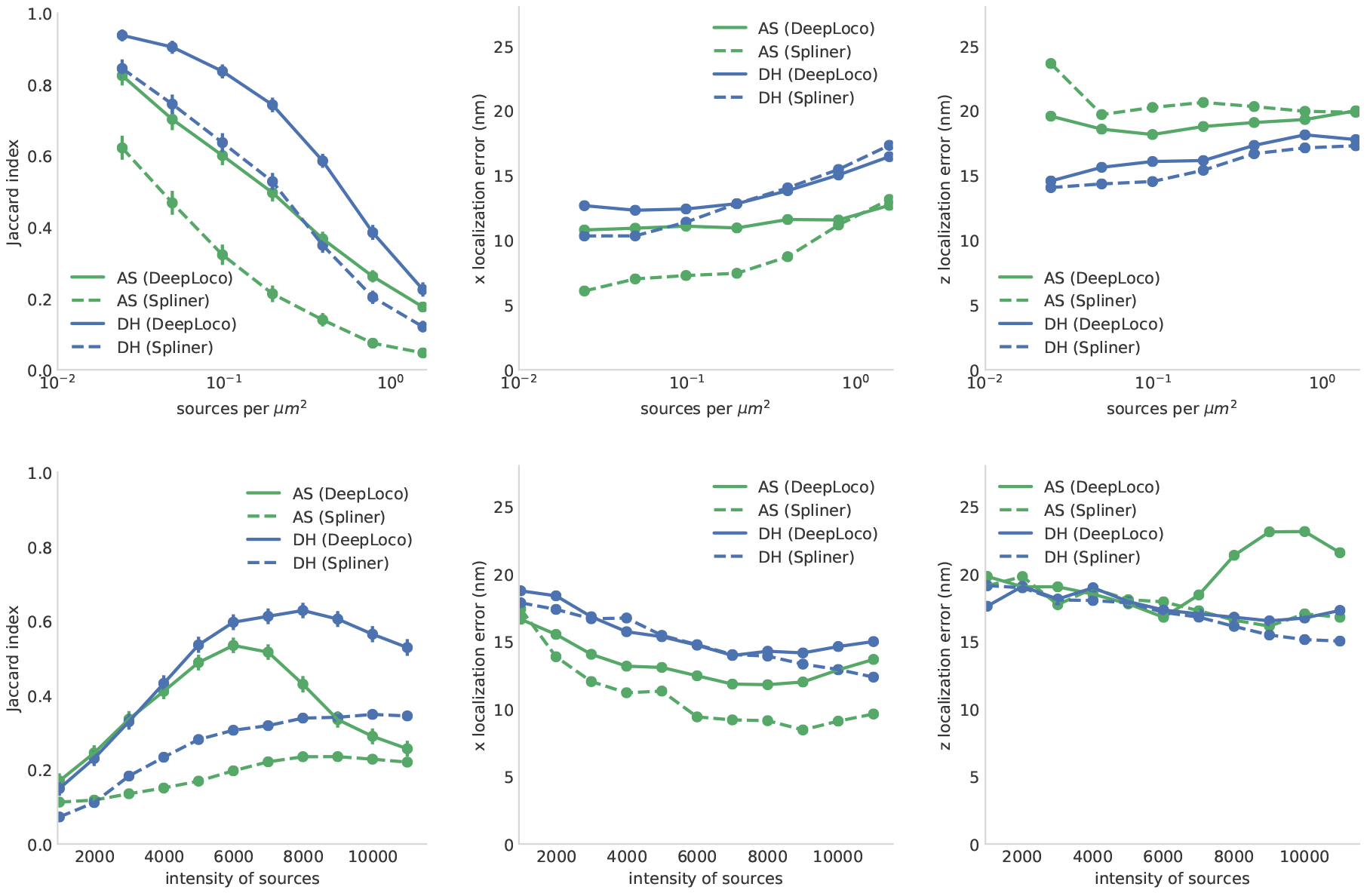
3D localization of synthetic data by DeepLoco and Spliner. All point matchings are computed with a 50nm tolerance radius. For experiments with varying source intensity we use a fixed density of 0.2 molecules per *μ*m^2^. Error bars reflect a 95% confidence interval on the mean estimated by the bootstrap. While the Jaccard index suggests that localization at very high densities (i.e. 10 sources per *μ*m) fails, both Spliner and DeepLoco produce reasonable images — suggesting that if the goal is to render an image the Jaccard index is not a representative metric.

**Table 1.** Reconstruction metrics for various algorithms across 3D datasets. All point matchings are computed with a 100nm tolerance radius. Values are mean ± standard deviation.

| Low Density SMLM 2016 3D Data |  |  |  |  |  |
| --- | --- | --- | --- | --- | --- |
| PSF | Algorithm | Jaccard | $x$ RMSE (nm) | $z$ RMSE (nm) | kernel loss |
| 3D-AS | DeepLoco | $0.83 \pm 0.33$ | $11.83 \pm 11.02$ | $24.80 \pm 17.85$ | $0.38 \pm 0.54$ |
| | Spliner | $0.77 \pm 0.37$ | $9.88 \pm 9.77$ | $22.34 \pm 18.45$ | $0.49 \pm 0.75$ |
| 3D-DH | DeepLoco | $0.84 \pm 0.30$ | $17.25 \pm 13.52$ | $22.35 \pm 16.71$ | $0.67 \pm 0.88$ |
| | Spliner | $0.74 \pm 0.36$ | $11.85 \pm 12.04$ | $15.99 \pm 15.04$ | $0.65 \pm 0.94$ |
| High Density SMLM 2016 3D Data |  |  |  |  |  |
| PSF | Algorithm | Jaccard | $x$ RMSE (nm) | $z$ RMSE (nm) | kernel loss |
| 3D-AS | DeepLoco | $0.51 \pm 0.16$ | $16.12 \pm 6.56$ | $27.30 \pm 8.56$ | $6.49 \pm 3.41$ |
| | Spliner | $0.37 \pm 0.16$ | $14.89 \pm 7.42$ | $27.83 \pm 10.91$ | $9.18 \pm 4.56$ |
| 3D-DH | DeepLoco | $0.39 \pm 0.19$ | $23.60 \pm 9.32$ | $28.89 \pm 9.98$ | $9.43 \pm 6.35$ |
| | Spliner | $0.27 \pm 0.14$ | $24.26 \pm 11.87$ | $25.13 \pm 11.93$ | $11.36 \pm 5.13$ |

**Figure. 4.**
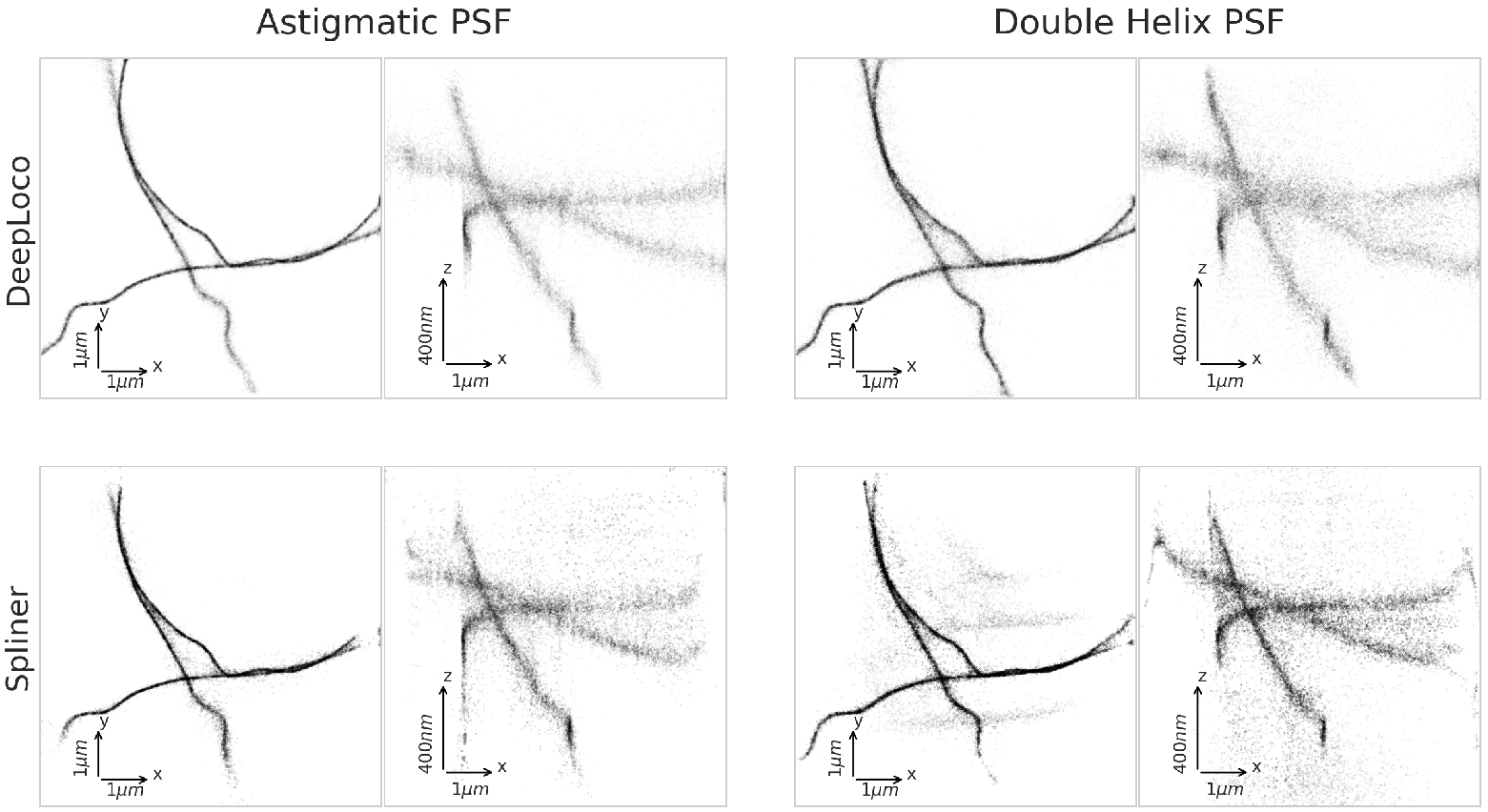
Rendered super-resolution images by DeepLoco and Spliner on the SMLM2016 MT0.N1.HD high-density dataset for both astigmatic and double-helix PSFs.

**Table 2.** Reconstruction metrics for DeepLoco and ADCG across high and low-density 2D datasets. All point matchings are computed with a 100nm tolerance radius.Values are mean ± standard deviation.

| Density | Algorithm | Jaccard | $x$ RMSE (nm) | kernel loss |
| --- | --- | --- | --- | --- |
| Low | DeepLoco | $0.89 \pm 0.26$ | $11.61 \pm 12.27$ | $0.22 \pm 0.45$ |
| | ADCG | $0.79 \pm 0.35$ | $14.95 \pm 13.27$ | $0.50 \pm 0.84$ |
| High | DeepLoco | $0.61 \pm 0.16$ | $18.22 \pm 7.24$ | $5.22 \pm 3.29$ |
| | ADCG | $0.53 \pm 0.15$ | $19.36 \pm 7.43$ | $6.63 \pm 3.60$ |

**Table 3.** Algorithm runtime for different datasets from the 2016 SMLM challenge

| PSF | Density | Algorithm (frame/sec) |  |  |
| --- | --- | --- | --- | --- |
|  |  | ADCG | Spliner | DeepLoco |
| 2D | low | 2.2 |  | 20859 |
|  | high | 0.44 |  | 20859 |
| 3D-AS | low |  | 40.6 | 20859 |
|  | high |  | 9.9 | 19742 |
| 3D-DH | low |  | 63.5 | 20859 |
|  | high |  | 9.3 | 20806 |

On high-density data, DeepLoco is roughly 40000 times faster than ADCG and 2000 times faster than Spliner.

## 7 Extensions and Variations

In this section we briefly describe some extensions to this work.

The successful application of machine learning techniques to the SMLM inverse problem opens up a new path to better SMLM algorithms: developing more accurate simulators. Raw SMLM data rarely looks like the output of the simulators we use in this paper. This discrepancy has two main causes: structured background fluorescence and aberrations in the optical system. In existing algorithms these issues are typically handled using delicate, heuristic preprocessing steps. Our approach suggests a different approach: training a neural network with a simulator that directly models background noise and aberrations.

One issue exposed in our experiments is the sensitivity of our networks to mismatch between the training distribution and the test distribution. For instance, and unlike any existing approach, our network can return *less* accurate localizations for extremely bright fluorophores. This bizarre behavior can be explained by a mismatch between the training distribution and the test distribution: networks trained to localize (relatively) low SNR sources do not generalize to higher SNR sources. Investigating and mitigating this issue would increase confidence in our approach.

From a statistical point of view, processing each frame independently is highly suboptimal: the photophysics of the fluorophore molecules is such that fluorophores active in one frame are often active in neighboring frames. Because localization accuracy is limited by the photon count of each emitter [31], analyzing multiple frames together could greatly increase accuracy. It’s also possible that analyzing multiple frames could help localizing much higher densities of emitters: at high densities emitters that are spatially overlapping still blink independently. Several existing techniques analyze multiple frames [4, 9, 44] but are limited by computational cost. The increased processing speed provided by using a feedforward neural network might allow efficient analysis of multiple frames and would be straightforward to implement.

Programmable spatial light modulators allow experimenters to change the point spread function of the microscope to, for instance, localize over much larger depths [41]. A neural network that takes the phase mask in addition to the raw image as input might be able to localize using a huge variety of point-spread functions.

A related question is how to reduce the expense of training the neural network for new experimental conditions (e.g. different noise levels or background conditions). While training time in our experiments was typically on the order of a few hours, preliminary experiments with warmstarting the optimization from pre-trained networks suggest this can be significantly reduced.

The loss function we introduce is a natural loss function for a variety of classification tasks where the labels are collections of parametric objects, for instance object detection [12]. Comparing the maximum mean discrepancy to existing loss functions would be interesting. Another possibility for SMLM or other classification problems with set-valued labels is to use different distances between discrete measures: for instance unbalanced optimal transport [6].

Finally, it is unclear how other machine learning techniques would do if applied to this problem. Indeed, it is quite possible that the machinery of deep learning is unnecessary and a simpler algorithm such as *k*-nearest neighbors would suffice.

## 8 Conclusion

A novel kernel-based loss function allowed us to train a neural network to directly localize sparse emitters in both two and three dimensions. DeepLoco is orders of magnitude faster than existing approaches, while achieving comparable accuracy. The success of DeepLoco suggests that there are regimes where coupling a naive black-box simulator with machine learning can efficiently solve inverse problems; even problems with complex, structured outputs. More physically accurate simulation, including accurately modeling phenomena that would preclude the application of traditional optimization-based approaches, could further improve accuracy.

## 9 Author Information

### 9.1 Author Contributions

EJ & NB conceived of original work, ran experiments, and wrote text. HB assisted in method evaluation and provided scientific feedback. Work was supervised by BR.

## Acknowledgements

The authors wish to thank Anna Thompson for help with Figure 1. We wish to thank Nick Antipa and Ren Ng for early discussions of this work.

This work was supported in part in part by DHS Award HSHQDC-16-3-00083, NSF CISE Expeditions Award CCF-1139158, DOE Award SN10040 DESC0012463, and DARPA XData Award FA8750-12-2-0331, and gifts from Amazon Web Services, Google, IBM, SAP, The Thomas and Stacey Siebel Foundation, Apple Inc., Arimo, Blue Goji, Bosch, Cisco, Cray, Cloudera, Ericsson, Facebook, Fujitsu, HP, Huawei, Intel, Microsoft, Mitre, Pivotal, Samsung, Schlumberger, Splunk, State Farm and VMware. B. Recht is supported by NSF award CCF-1359814, ONR awards N00014-14-1-0024 and N00014-17-1-2191, the DARPA Fundamental Limits of Learning (Fun LoL) Program, a Sloan Research Fellowship, and a Google Faculty Award. N. Boyd was funded by a Google Hertz Fellowship. E. Jonas is supported by ONR award N00014- 17-1-2401. H. Babcock is supported by the Center for Advanced Imaging at Harvard University

## Footnotes

1 Or, in some cases, an approximation to the full posterior.

2 Note that each of these algorithms was developed by different authors on this paper

